# Diffeomorphic registration with intensity transformation and missing data: Application to 3D digital pathology of Alzheimer’s disease

**DOI:** 10.1101/494005

**Authors:** Daniel Tward, Timothy Brown, Yusuke Kageyama, Jaymin Patel, Zhipeng Hou, Susumu Mori, Marilyn Albert, Juan Troncoso, Michael Miller

## Abstract

This paper examines the problem of diffeomorphic image mapping in the presence of differing image intensity profiles and missing data. Our motivation comes from the problem of aligning 3D brain MRI with 100 micron isotropic resolution, to histology sections with 1 micron in plane resolution. Multiple stains, as well as damaged, folded, or missing tissue are common in this situation. We overcome these challenges by introducing two new concepts. Cross modality image matching is achieved by jointly estimating polynomial transformations of the atlas intensity, together with pose and deformation parameters. Missing data is accommodated via a multiple atlas selection procedure where several atlases may be of homogeneous intensity and correspond to “background” or “artifact”. The two concepts are combined within an Expectation Maximization algorithm, where atlas selection posteriors and deformation parameters are updated iteratively, and polynomial coefficients are computed in closed form. We show results for 3D reconstruction of digital pathology and MRI in standard atlas coordinates. In conjunction with convolutional neural networks, we quantify the 3D density distribution of tauopathy throughout the medial temporal lobe of an Alzheimer’s disease postmortem specimen.

**Author summary:** Our work in Alzheimer’s disease (AD) is attempting to connect histopathology at autopsy and longitudinal clinical magnetic resonance imaging (MRI), combining the strengths of each modality in a common coordinate system. We are bridging this gap by using post mortem high resolution MRI to reconstruct digital pathology in 3D. This image registration problem is challenging because it combines images from different modalities in the presence of missing tissue and artifacts. We overcome this challenge by developing a new registration technique that simultaneously classifies each pixel as “good data” / “missing tissue” / “artifact”, learns a contrast transformation between modalities, and computes deformation parameters. We name this technique “(D)eformable (R)egistration and (I)ntensity (T)ransformation with (M)issing (D)ata”, pronounced as “Dr. It, M.D.”. In conjunction with convolutional neural networks, we use this technique to map the three dimensional distribution of tau tangles in the medial temporal lobe of an AD postmortem specimen.

## Introduction

High throughput neuroinformatics is emerging in neuroscience [1, 2]. Atlas based image analysis plays a key role, as it enables information encoded by millions of independent voxel measurements to be reconstructed in ontologies of the roughly 100 evolutionarily stable structures. At the 1 millimeter scale there are many atlases, including Tailarach coordinates [3], Montreal Neurological Institute (MNI) [4], and Mori’s diffusion tensor imaging (DTI) white matter atlases [5], which define the locations of neuroanatomical structures as well as important structural and functional properties such as volume, shape, blood oxygen-level dependent (BOLD) signals, etc. At micron and meso-scales there are several atlases including Mori’s and Allen brain atlas [6, 7] with their associated region and cell-types.

High-throughput image analysis relies on some form of dense brain mapping. At the millimeter scale there have been several approaches to computational anatomy with deformable templates [8–13], atlas estimation [14, 15], and applications in white matter [16] or even cardiac imaging [17, 18]. Many of these algorithms have been extended to micron scales, such as for CLARITY [19–21]. However, most of these datasets and the methods used to analyze them are based on high quality clinical images. At the frontier of neuroscience are images that are less controlled. In this work we focus on digital pathology, where micron thick tissue slices are prone to damage and subsets which are completely missing, and where a host of stains can result in a multitude of different image intensities. To bring digital pathology to the era of high throughput neuroinformatics, brain mapping algorithms need to be expanded to handle missing data and multiple modalities with perhaps radically different contrast profiles.

The community has developed a host of techniques for addressing these situations partially. Image masking allows algorithms to ignore missing or unreliable data, and similarity functions such as normalized cross correlation [22, 23] or mutual information [24] allow mapping between different image contrasts to an extent. These tools are available in standard software packages such as the Insight Toolkit (ITK) [25]. However, modern image mapping techniques such as multi atlas [26] or Bayesian segmentation [27] require not just similarity functions, but a complete generative model that describes the likelihood of each observed image voxel.

In this work we develop a generative probabilistic model that accounts for differences in contrast and missing or censored data. We use the random orbit model of Computational Anatomy [9, 28] in which the space of histological images is an orbit of exemplar templates under smooth coordinate transformation, as well as transformation of contrast. The models for coordinate transformations are taken from diffeomorphometry [29, 30]. We wish to accommodate differing contrasts associated to different histological staining such tau, amyloid, myelin, and a variety of contrasts such as Nissl and fluorescence at the meso- and micro-scale. To achieve this, we model the space of contrasts manifest in the orbit of observed images as polynomial functions of the template. In the case of first order polynomials, this gives affine models which we show reduce to the normalized cross-correlation cost function. Application of higher order polynomials can describe non-monotone transformations which swap the order of intensities. We name this mapping framework, “(D)eformable (R)egistration and (I)ntensity (T)ransformation”, pronounced as “Dr. It”.

To accommodate effects such as folded or missing tissue, we include additional homogeneous atlases, and model each pixel in an observed image as a realization of one transformed atlas from this family. Pixels corresponding to missing tissue, data deletion, or global censoring are interpreted predominantly via the “background only” atlas. Pixels corresponding to anomalous intensities are interpreted predominantly via the “artifact” atlas. The unknown atlas label at each pixel is interpreted as missing data, and its conditional mean is estimated using the Expectation Maximization algorithm [31]. The conditional mean is interpreted as the posterior probability of a particular atlas in the multi atlas random orbit model. We name the version of our algorithm that includes missing data “(D)eformable (R)egistration and (I)ntensity (T)ransformation with (M)issing (D)ata”, pronounced as “Dr. It, M.D.”.

In this paper, our framework is studied using simulated examples, and those from digital pathology and MRI. We apply these techniques to an important application in Alzheimer’s disease (AD), computing the 3D density of tau neurofibrillary tangles, a key pathologic feature of AD. Tangles are detected from histology images using convolutional neural networks, registered to ex vivo MRI, and mapped to the standard coordinates of the Mai Paxinos Voss atlas [32].

## Methods

### The contrast transformation problem

Our generative model for the space of observables or target images builds upon the deformable templates of [28, 33]. Observed images are transformations of collections of exemplars or atlas images 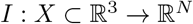, which may be single valued (*N* = 1) such as a T1 MRI, or multi-valued such as red green blue (RGB, *N* = 3). The atlases experience two kinds of transformations: deformations on the background space *φ*: *X* → *X*, and transformations of the image intensity 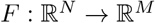. The mapping *F* depends on imaging instrumentation and tissue properties. Considering *N* ≠ *M* can accommodate mappings between many different modalities.

We define the deformations of coordinates as diffeomorphisms, *φ* ∈ *Diff*, generated using flows as in [9], *φ* = *θ*_1_, 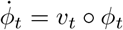, *t* ∈ [0, 1], *φ*_0_ = *id, v*_*t*_ the Eulerian vector fields of the flow. The mapping of image intensity is defined parametrically, 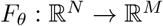, *θ* ∈ Θ unknown in some parameter set. In this work we consider *F* as polynomials with Θ the set of coefficients.

The random imaging model takes the observable images *J*(*x*), *x* ∈ *X* as a conditionally Gaussian random field having mean given by the transformed template, and white noise variance *σ*^2^:

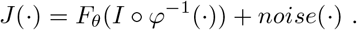

The log-likelihood, as a function of the unknown parameters is

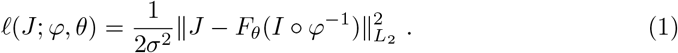

While the dimension of *θ* is finite, that of the diffeomorphism is infinite. We therefore use penalized methods, introducing a Sobolev norm on the vector fields 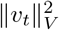 as a running penalty. We force *v* ∈ *V* to be a reproducing kernel Hilbert space with kernel *K* defined through the differential operator *A*: *v* ∈ *V* ↦ *Av* ∈ *V*\*, *V*\* the dual-space of smooth vector fields *V*. Then the norm written as generalized function integration becomes

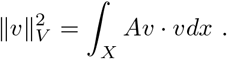

The penalized likelihood becomes for 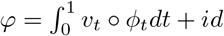, and 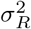 a regularization parameter:

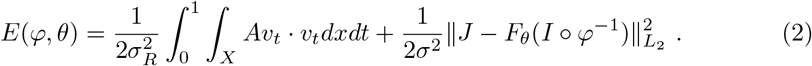

To calculate the penalized maximum likelihood estimators we assume the mapping *F*_*θ*_ differentiable with 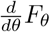 an *M × B* vector, *B* being the number of basis functions in our polynomial.

#### Theorem 1.

Minimizers of he penalized likelihood of (2) satisfy

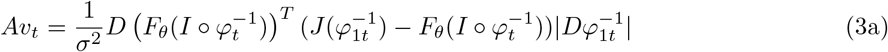

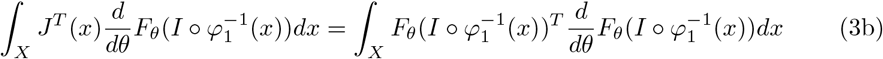

The first equation (3a) is the necessary condition for the stationary solution with respect to the deformation controlled by the vector field, the original LDDMM equation of Beg [34], where 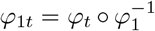 is a mapping from time 1 to time *t*. The equation (3a) is computed by application of the chain rule. Define for notational convenience *Ĩ* ≐ *I* ○ *φ*^−1^. First the derivative of *E* with respect to *F*_*θ*_(*Ĩ*), second *F*_*θ*_(*Ĩ*) with respect to *Ĩ*, and third *Ĩ* with respect to the deformation field. The first and second steps are combined as

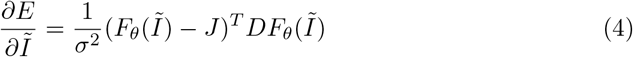

The third step is discussed in [34]. This equation can be solved using a standard gradient descent approach. The equation (3b) is the necessary condition with respect to the contrast or photometric parameters. Here we consider *F*_*θ*_ as a linear combination of *B* polynomial basis functions, and so (3b) is a linear system solved exactly at each iteration of gradient descent.

Minimizing over both *θ* and transformation parameters means the result of registration will be independent of the family of transformations indexed by *θ*. Alternatively, we can consider minimizing over *θ* first, leading to an invariant cost function

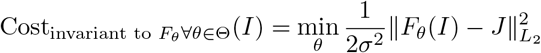

#### Corollary

(Unknown Affine Transformations Correspond to Normalized Cross Correlation Squared). When *F* is an affine map from 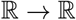, our registration results will be invariant to affine transformations of our atlas, and (3b) can be solved analytically. For *N* = *M* = 1 and 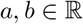, *F_a,b_*(*t*) = at + *b*, minimizing 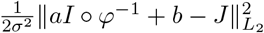 is equivalent to maximizing the normalized cross correlation squared of *I* ○ *φ*^−1^ with *J*.

*Proof*. For any fixed *φ*, optimal values of *a, b* can be found via a standard linear least squares estimation result, which gives *a* = Cov(*Ĩ, J*)*/* Var(*Ĩ*) and 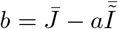 where 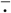 corresponds to the expected value and expectation is taken by averaging over all voxels in the images.

Plugging these into our *L*_2_ cost gives

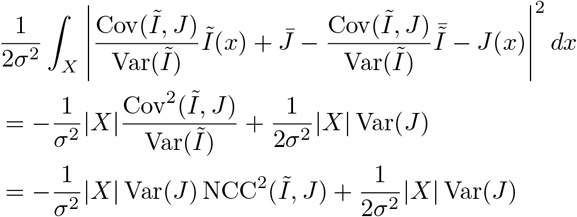

Up to constants that do not depend on the deformation, minimizing sum of square error with an unknown affine intensity transformation is equivalent to maximizing normalized cross correlation (NCC) squared.

### The Missing Data Problem

We approach the problem of missing or censored data using the Expectation Maximization algorithm [31]. We discretize our problem by associating to the images the lattice of sites {Δ*x*_*i*_, *i* = 1,…, *S*} which are a disjoint partition (voxels) 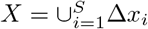. Each atlas type 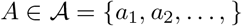 includes a deformation given by 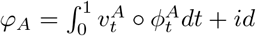, so the discrete values can be written as

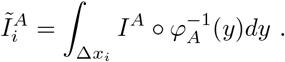

The observable random field *J*_*i*_, *i* = 1,…, *S*, is conditionally Gaussian with constant variances 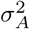 and mean fields 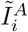 determined by the atlas type *A*. In practice, we choose *I*^*a*1^ as our atlas image, and *I*^*ai*^ for *i* ≠ 1 to be constant images. Since they are constant, we need not optimize over the deformations 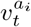.

Associate to the measured *incomplete-data* **Y** = {*J*_*i*_, *i* = 1, 2,…, *S*} the *complete-data* **X** = {(*J*_*i*_, *A*_*i*_), *i* = 1,…, *S*}, an augmentation with labels determining the atlas types 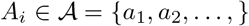. Model *J*_*i*_ as a Gaussian random variable with mean 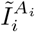 and variance 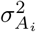. The complete-data penalized log-likelihood becomes:

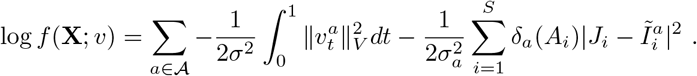

The Kronecker-delta *δ*_*a*_(·) is 1 when the argument is *a*, and zero otherwise. The Expectation step (E-step) replaces these functions with their expected value, a posterior probability *π*_*i*_(*a*) at each voxel.

#### Theorem 2.

The Expectation Maximization algorithm performs:

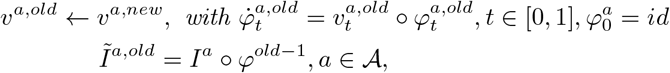

*E-step: E logf*(**X**; *v*)*|***Y**, *v*^*a,old*^

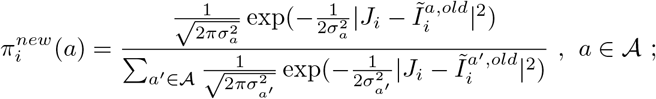
*M-Step:* arg max *E* log *f*(**X**; *v*)*|***Y**, *v*^*old*^

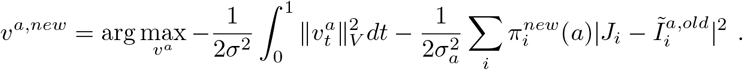

Iterations *v^a,new^* with 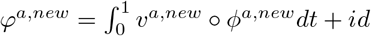 increase in incomplete-data log-likelihood.

The M step updates *v*_*t*_ which is just weighted LDDMM with a weighted L2 cost. Equation (3b) is updated to include the posterior weights (derived for example in [35]), and (3a) is updated as weighted least squares, giving

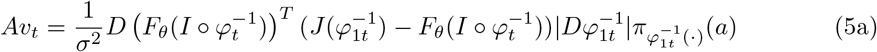

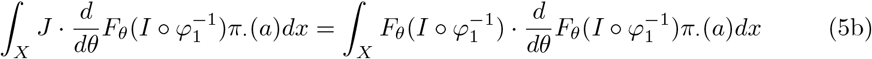

In (5a) and (5b), *π*.(*a*) are considered functions of space rather than voxel indices.

### Post mortem imaging

Preparation and scanning of brain tissue was performed by the neuropathological team at the Johns Hopkins Brain Resource Center (BRC) and the laboratory of Dr. Susumu Mori. The specimen was a 1290 gram brain from a 93 year old male, with a clinical diagnosis of Alzheimer’s disease dementia. The autopsy diagnoses included: Alzheimer’s disease neuropathologic change, high level (A3,B3, C2) [36]; CERAD neuritic plaque score B [37]; neurofibrillary Braak stage VI/VI [38]; subacute infarcts frontal, temporal, and basal ganglia; old infarct of pons; with clinical-pathological comment: “Mixed dementia, AD and vascular. The AD component appears to predominate”. The fixed brain tissue was divided into six coronal blocks of the temporal lobe that contain the entorhinal cortex, the hippocampus, and the amygdala. The orientation of the blocks correspond as closely as possible to the coordinate system of the Mai Atlas. Each block of brain tissue was scanned with a high field 11.7T MRI scanner.

The nuclear magnetic resonance (NMR) sequence was based on a 3D multiple echo sequence [39, 40] with four echoes acquired for each excitation. The diffusion-weighted images were acquired with a field of view of typically 40 × 30 × 16 mm and an imaging matrix of 160 × 120 × 64, which was zero-filled to 320 × 240 × 128 after the spectral data were apodized by a 10 percent trapezoidal function. The pixel size was native 250 micron isotropic. Eight diffusion-weighted images were acquired with different diffusion gradient directions, with b-values in the 1,200 - 1,700 s/*mm*^2^ range. For diffusion-weighted images, a repetition time of 0.9 s, an echo time of 37 ms, and two signal averages were used, for a total imaging time of 24 hours.

The MRI scanning procedure resulted in several distinct images which must be themselves aligned after imaging. For this we developed an interactive tool for visualizing and transforming each imaged block to match with the others, and for rigidly positioning the aligned blocks in the Mai Atlas coordinate system. Shown in Fig. 1 left is an example of our images and their alignment. The aligned blocks were labeled by a neuroanatomist for relevant medial temporal lobe structures (entorhinal cortex, subiculum, Cornu Ammonis (CA) fields, compartments of dentate gyrus, alveus). These labels are shown as a surface reconstruction in Fig 1 right. The superimposed lines correspond to the rostral-caudal pages of the Mai-Paxinos-Voss atlas (z axis), with their 1cm grid lines along the x-y axes of each page. Our entorhinal cortex definition (orange) includes the medial bank of the collateral sulcus. This sulcal region [41], also referred to as the trans entorhinal cortex, corresponds to the earliest location of AD pathology accumulation visible at autopsy [42]. Atrophy in this region has been detected at the population level in subjects with mild cognitive impairment [43] before other changes are visible [44].

**Fig 1.**
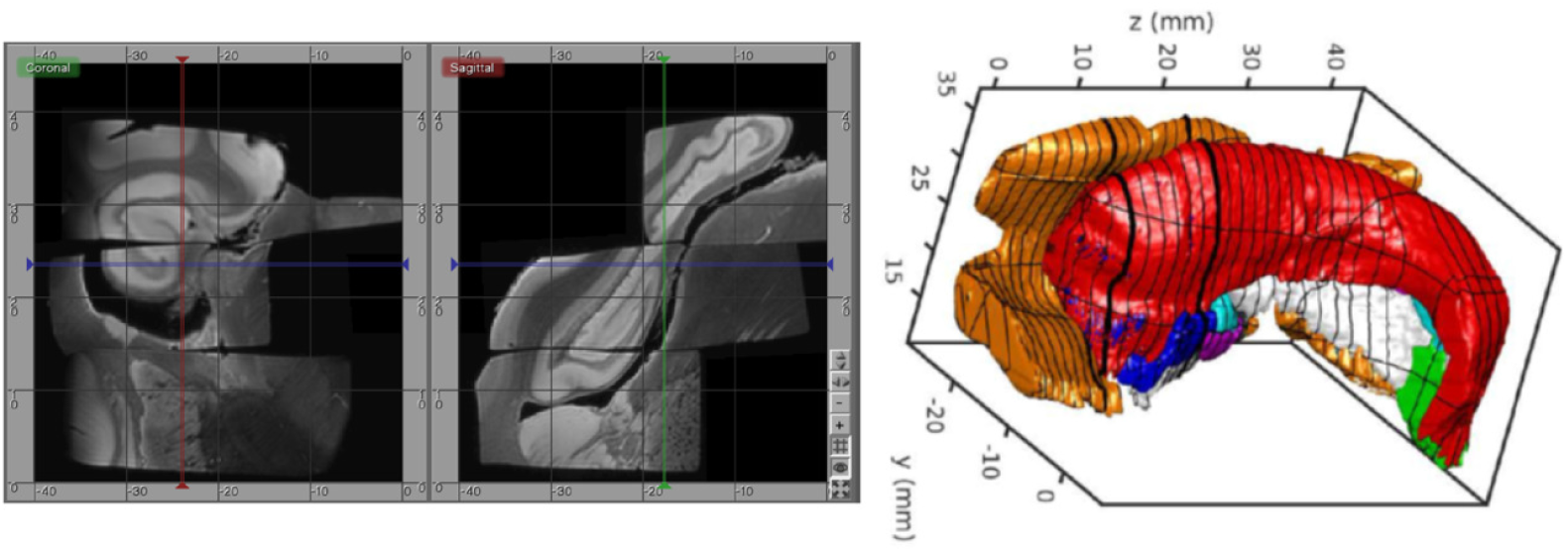
High-field MTL volume from MR. Left: MR images of tissue blocks aligned in 3D. Right: surface rendering of manual segmentations with Mai Atlas coordinate system superimposed.

After the tissue blocks underwent high field imaging, they were sectioned for histological examination. The tissue was sectioned sectioned at 200*μ*m intervals: 20 slices of 10-*μ*m thickness and 5 slices of 40-*μ*m thickness. The thin-sliced sections are prepared with stains focused on AD pathology: Nissl, silver (Hirano method), Luxol fast blue (LFB) for myelin, and immunostained for A*β* (mab 6E10) and tau (PHF1). Several examples of our tau, amyloid, and myelin stained sections are shown in the result section, Fig. 4 to 9. For future analysis, these stains are complemented by additional stains to examine the tissue for related comorbidities and other neurodegenerative disorders, such as lewy body disease and frontotemporal dementia, including: immunostains for *α*-synuclein, ubiquitin, TDP-43, GFAP (astrocytes), Iba1 and CD68 (microglia), collagen IV (blood vessels) and reelin (entorhinal cortex layer II neuronal protein) 13. The thick-sliced sections will be used for quantitative cell and neuron counts and density studies of dendritic and synaptic markers.

**Fig 9.**
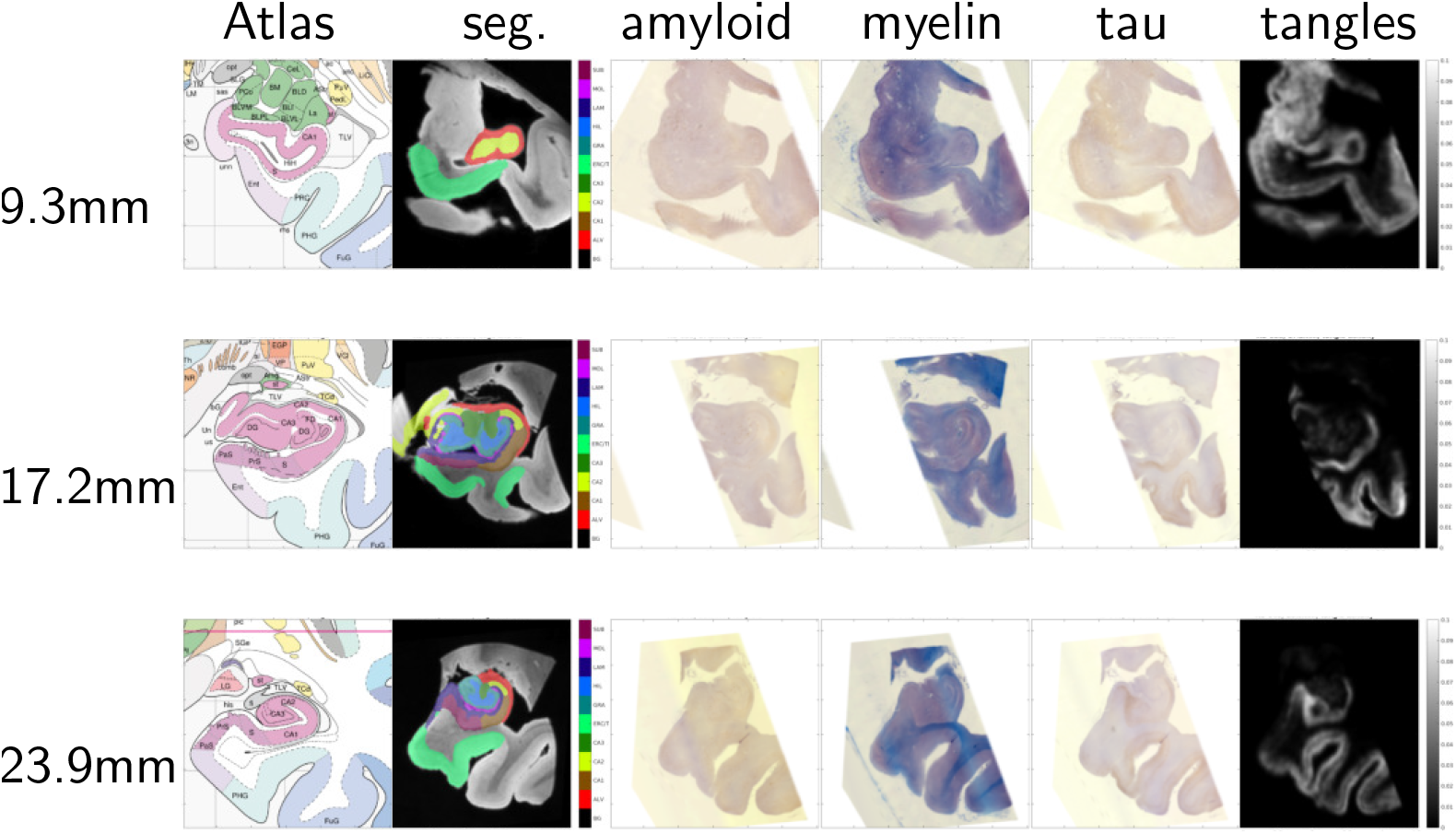
Mapping MRI sections to Mai-Paxinos sections. High-field atlas section at 9.30 mm (top row), 17.20 mm (middle row) and 23.90 mm (bottom row) along caudal-rostral axis of histology sections in Mai-Paxinos coordinates.

### Image mapping experiments

Below we present several 2D experiments to demonstrate the applicability and verify the validity of our method. In each case, rigid and deformable registration is performed simultaneously using gradient descent, and deformable registration is implemented using LDDMM (3a) or weighted LDDMM (5a). For registration purposes, histology images are downsampled by averaging over a 32 × 32 window

Our final example is to demonstrate an important application in a 3D histology pipeline, quantifying the 3D distribution of tau tangles in Alzheimer’s disease. Here we work with a sequence of transformations between 3D post mortem MRI and 2D histology slices.

1. 3D deformation
2. 3D rigid positioning
3. Estimation of slice spacing (*z* axis scale)
4. Estimation of pixel size (*x, y* axis uniform scale)
5. 2D rigid motion (for each slice and stain)
6. Cubic intensity transformation (for each slice and stain)

Rigid, scale, and deformation parameters are all jointly optimized using gradient descent. Deformable registration uses a relatively small gradient descent step size allow linear transformations to be close to optimal at all times.

This pipeline quantifies the distribution of tau tangles in each 2D slice using a convolutional neural network in tensorflow [45]. Input data is 56 × 56 regions of interest. The network uses 3 convolution layers with a 5 × 5 kernels, max pool downsampling by a factor of 2 × 2, followed by a fully connected layer and a cross entropy loss function. The center pixel of each region is classified as belonging to one of 3 classes “tau tangles”, “other tissue”, or “background”. 2391 training examples were used, with 8.3% positive, and neural network weights were trained using the Adam optimizer [46]. Every pixel in our histology data was classified using a sliding window approach. After mapping this data into the coordinates of the Mai atlas, we report the total area of tau tangles within the entorhinal cortex, subiculum, and CA1-3 for each atlas page. To place these numbers in context, we also report the total area of these structures (which may be affected by missing tissue), and the fraction of this area covered by tau tangles.

## Results

### Mapping simulated images with artifact and missing data

To demonstrate the method we start with simulated images. Fig 2 shows the atlas (top left panel) and target (top center panel). Contrast is chosen so that the atlas is appears like a T1 MR brain image (darker gray matter and brighter white matter) and the target appears like a T2 MR brain image. The target also contains a bright streak artifact and missing tissue. Specifically, the atlas background has intensity 0, gray matter 1, and white matter 1.25. The target background has intensity 0, gray matter 0.9, white matter 0.675, and artifact 5. Both images have additive white Gaussian noise with standard deviation 0.05, and are blurred with a Gaussian kernel with standard deviation 2/3 pixels over a 5 × 5 pixel window.

**Fig 2.**
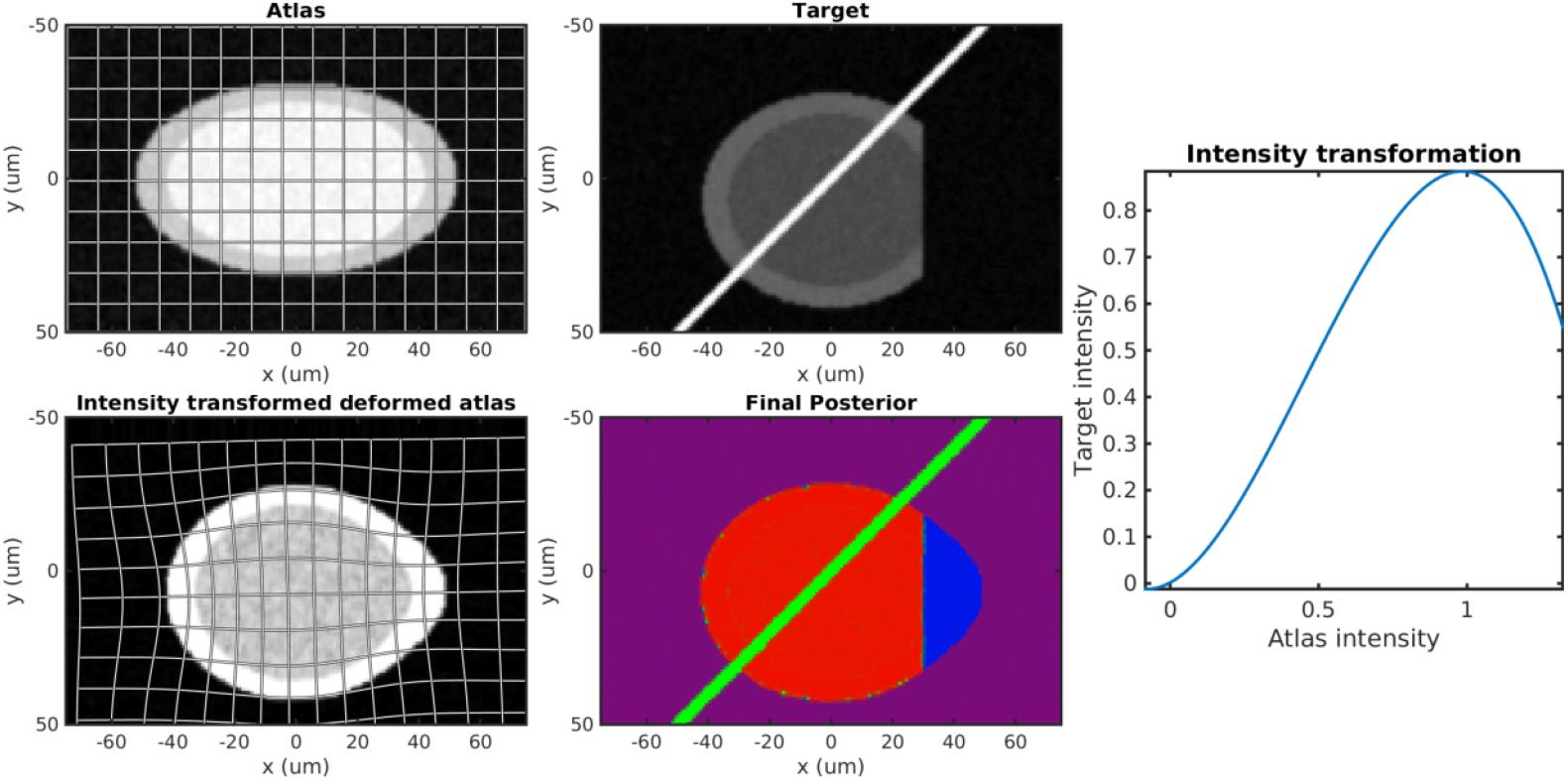
Simulation results. Top row: atlas (left) and target (right). Bottom row: Transformed atlas to target; estimated final weights; estimated non-monotonic polynomial *F*.

The bottom left panel of Fig. 2 shows that a cubic intensity transformation (right panel) is sufficient to permute the order of gray and white mater, allowing for accurate matching of the cortical boundary. The bottom center panel shows the posterior probabilities of the three atlases shown as components of an RGB image. The pixels are correctly classified with the atlas image in red, artifact in green, and missing tissue in blue. Note that the background is magenta because “atlas image” and “missing tissue” both describe the image intensity equally well.

Figure 3 shows the failed results of existing mapping methods, using a linear contrast transform only (i.e. normalized cross correlation), and a fixed mask (top row) or no mask (bottom row). With a fixed mask, the artifact is still handled appropriately. An inversion of contrast is estimated, which is appropriate within the masked region. However, missing tissue is not distinguished from normal background, so the informative “cortex/background” boundary is treated equivalently to the uninformative “cut tissue” boundary, resulting in very poor alignment. With no mask, huge distortions in shape occur as the atlas is squeezed to match the shape of the target with missing tissue, and stretched to follow the bright artifact.

**Fig 3.**
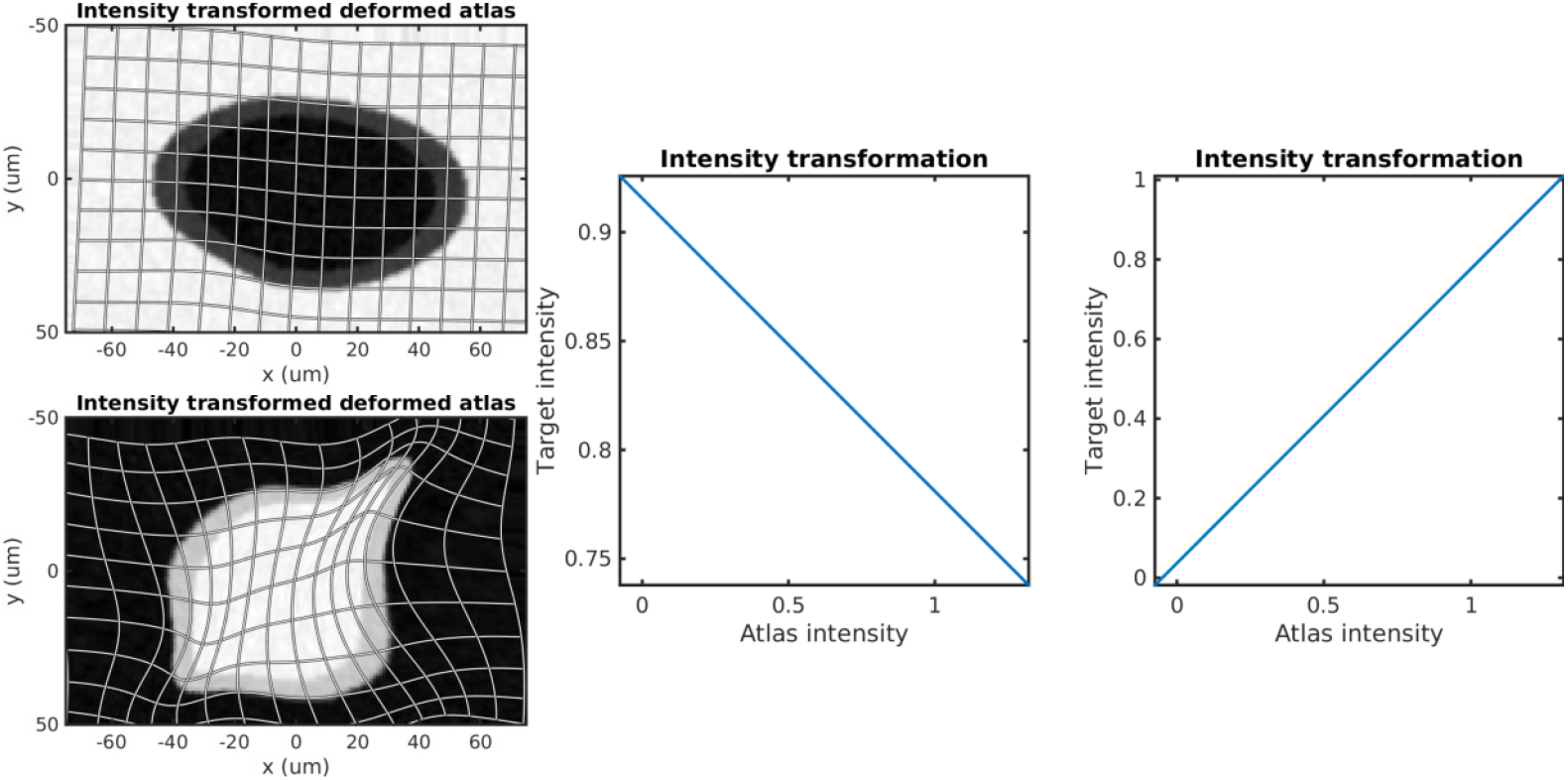
Alternative methods. Top left shows the results using a fixed binary mask, with center showing the estimated monotonic intensity transformation. Bottom left shows the result using no mask, with right showing the estimated intensity transformation.

**Fig 4.**
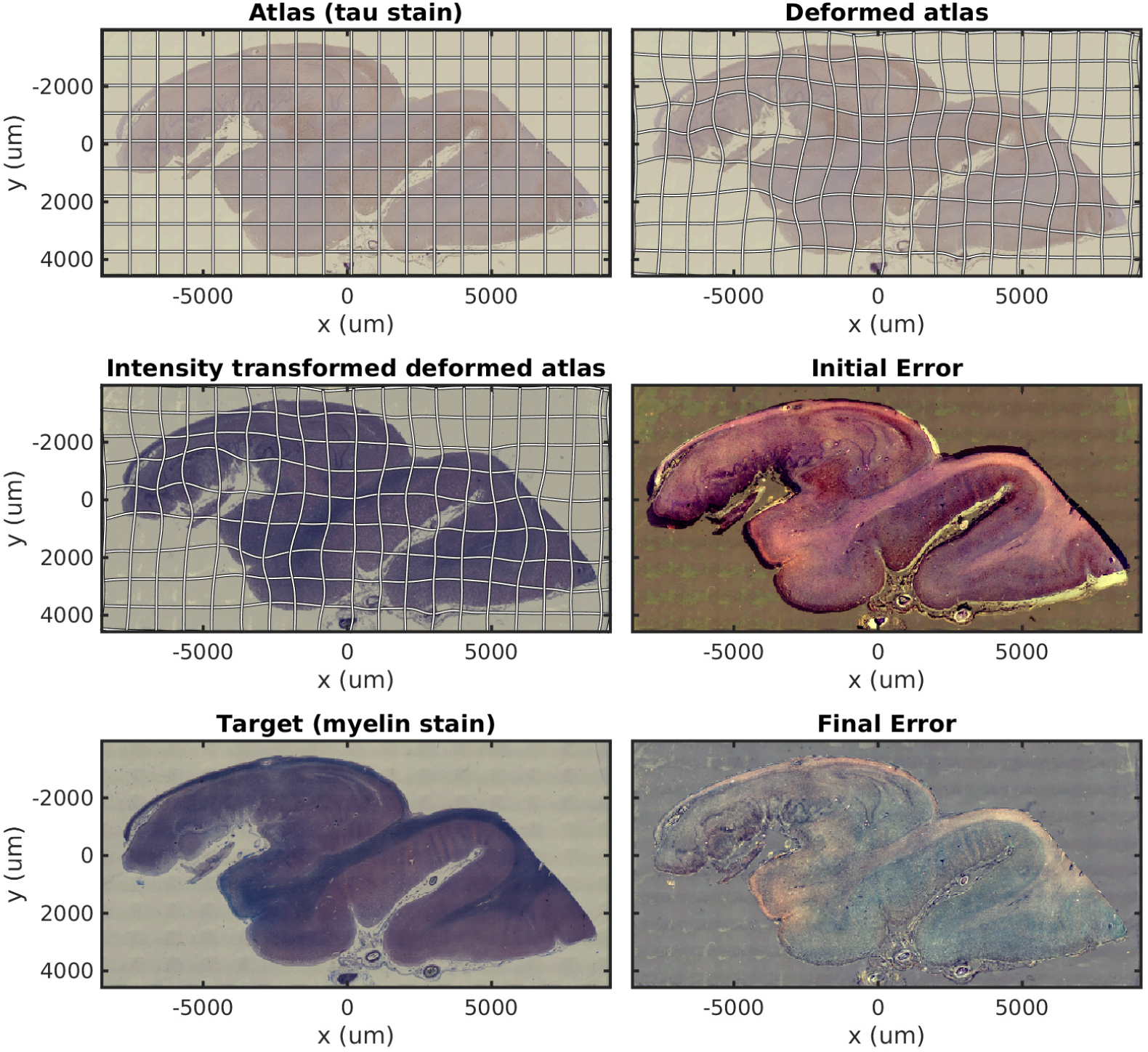
Mapping across modality from tau to myelin. A tau stained section through the hippocampus is mapped to a neighboring myelin stained section using a cubic polynomial intensity transform.

### Mapping histology with missing data and different stains

In Fig. 4 we show results mapping a tau stained section of the medial temporal lobe to an immediately adjacent section stained with LFB. We perform intensity transformation using a nonmonotonic cubic polynomial, allowing for a swapping of brightness from gray matter (1) → white mater (2) → background (3) in tau, to (2) → (1) → (3) in LFB. As a map from 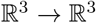, this corresponds to 60 unknown parameters (1 constant, 3 linear, 6 quadratic, 10 cubic, for each of 3 dimensions). This example illustrates the intensity transformation component of our algorithm in isolation.

In Fig. 5 we map a tau stained slice of medial temporal lobe to its neighbor which has significant missing data due to damaged tissue. This illustrates the missing data component of our algorithm in isolation.

**Fig 5.**
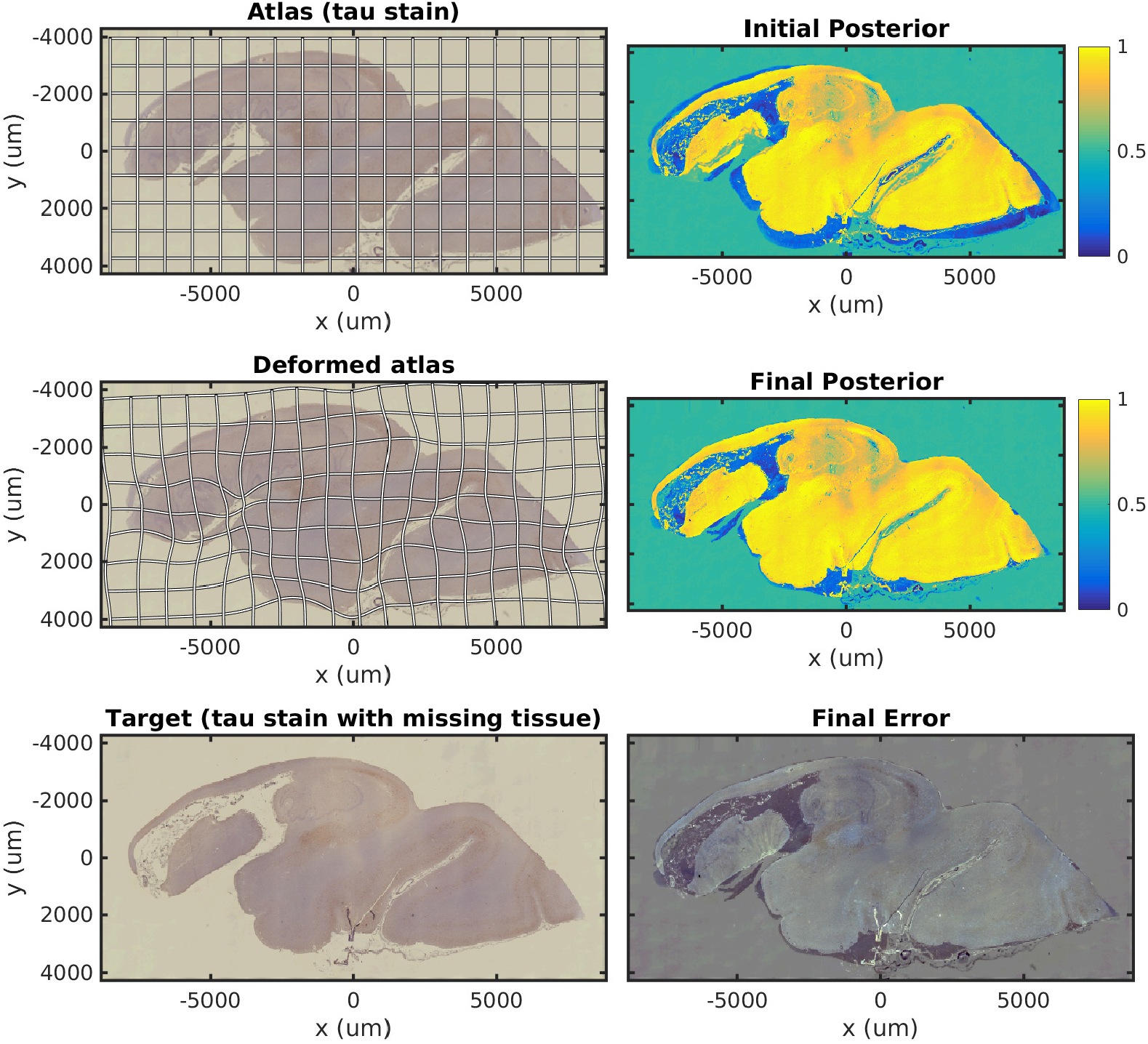
Mapping a tau section with missing tissue. A tau stained section through the hippocampus is mapped to a neighboring damaged section. The weights shown correspond to posterior probability that a given pixel corresponds to our atlas image, with high probability in yellow, probability 0.5 in green, and low probability in blue.

In Fig. 6 we show results mapping a tau stained section of the medial temporal lobe to an adjacent damaged slice stained with LFB. This illustrates the intensity mapping and missing data components of our algorithm simultaneously.

**Fig 6.**
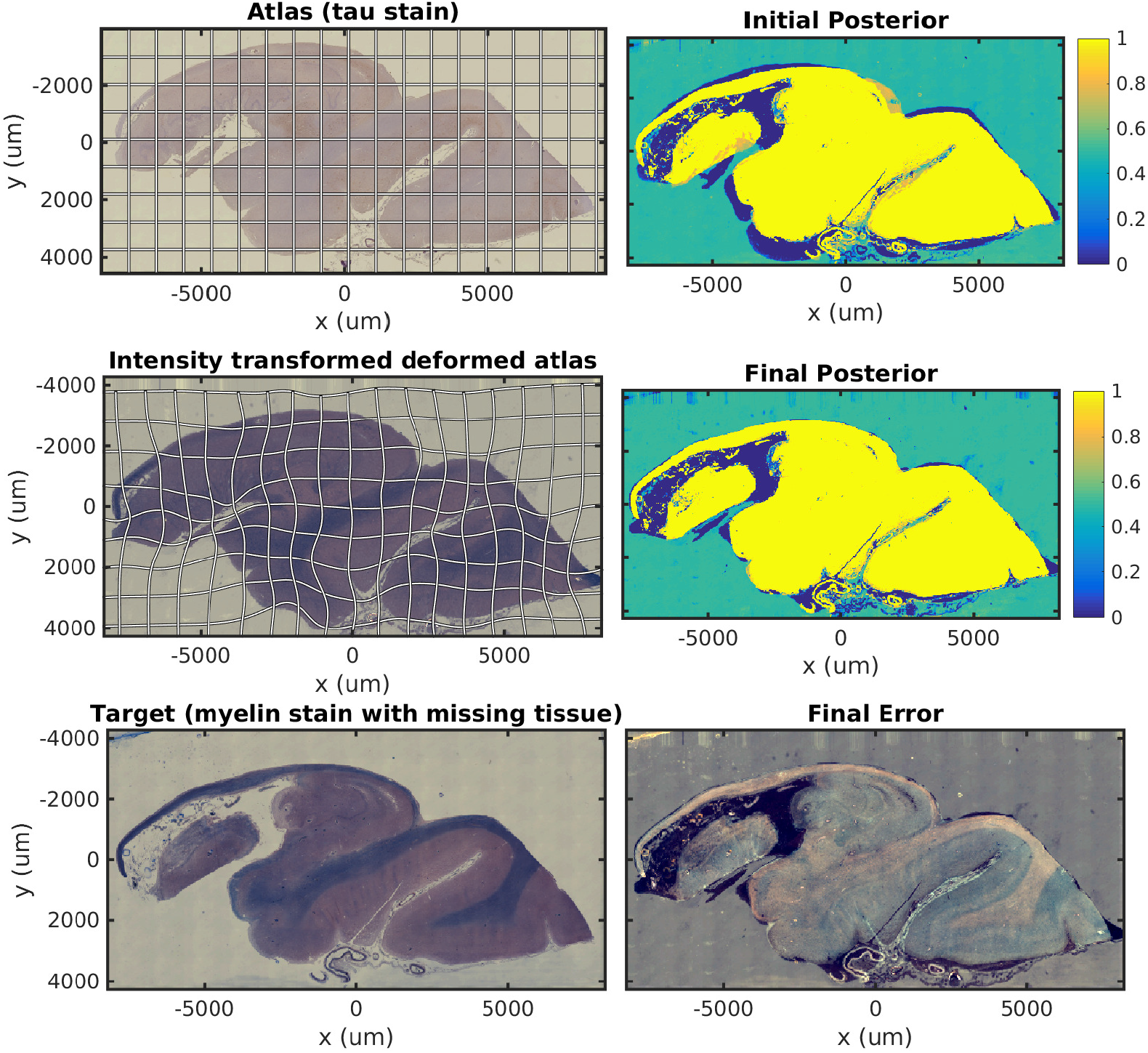
Mapping across modality from tau to myelin with missing tissue. A a tau stained section through the hippocampus is mapped to a neighboring LFB stained damaged section. The weights shown correspond to posterior probability that a given pixel corresponds to our atlas image, with high probability in yellow, probability 0.5 in green, and low probability in blue.

### Mapping histology data to Mai atlas coordinates

Alignment between 3D post mortem MRI, and each of our three 2D stains are shown in Fig 7. For each stain, we show histology images, intensity transformed MRI, and original aligned MRI. The LFB stain in particular shows significant variation in contrast profiles from slice to slice, which is handled effectively by our method. All intensity transformations use a cubic polynomial for each of the red, green, and blue channels on each slice, which corresponds to 12 parameters.

**Fig 7.**
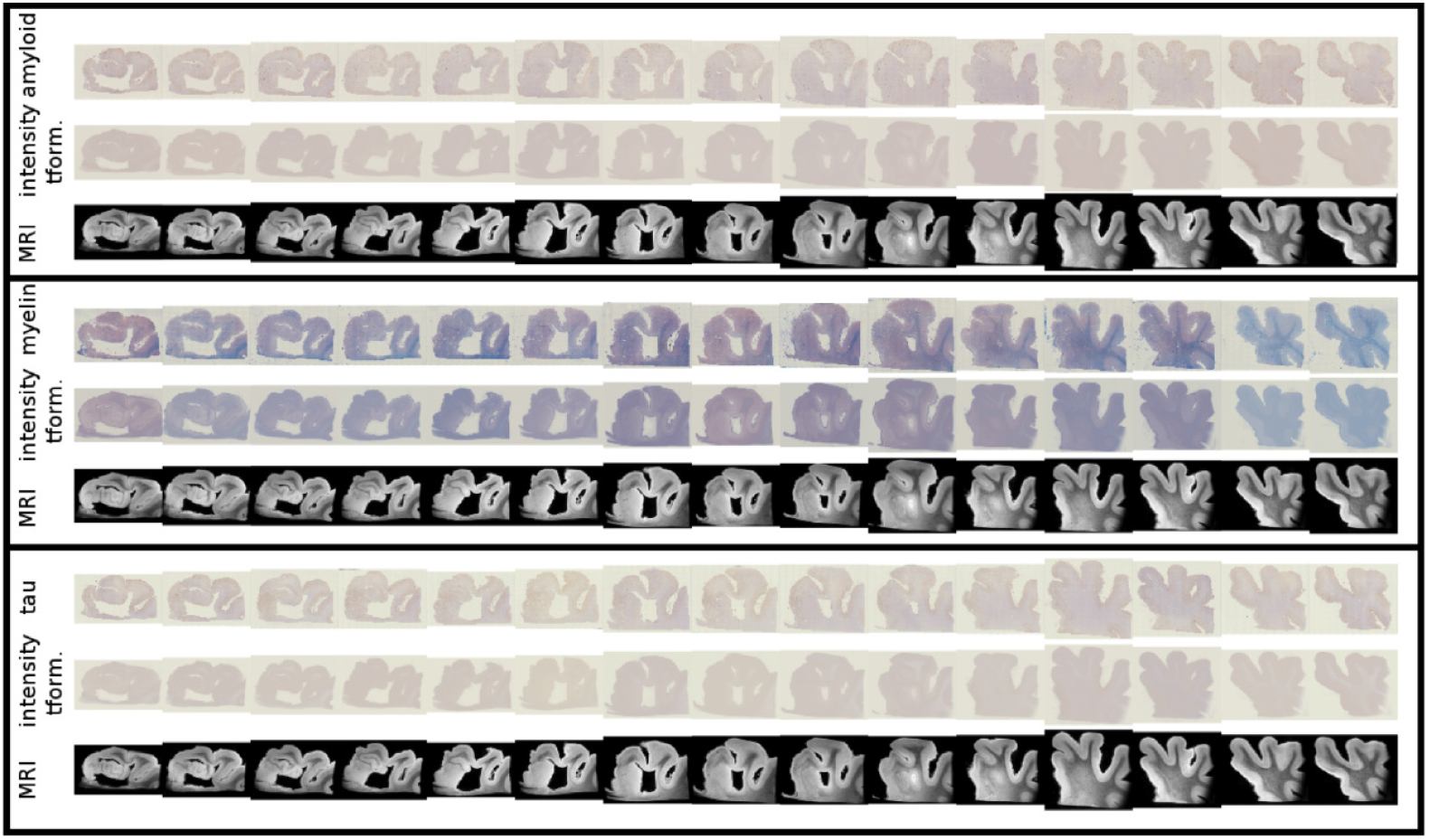
Mapping from MRI to three histology slices. Alignment between our 3D MRI and histology slices at 15 locations with three stains is shown. Top to bottom: amyloid, MRI intensity mapped to amyloid, aligned MRI, LFB, MRI intensity mapped to LFB, aligned MRI, Tau, MRI intensity mapped to tau, aligned MRI.

Example results of our tau detection algorithm are shown in Fig 8. We achieve an accuracy of 0.996 on a test set of 500 examples. Detection results are shown at 3 different scales, with individual detections at the micron level, and densities at the millimeter level.

**Fig 8.**
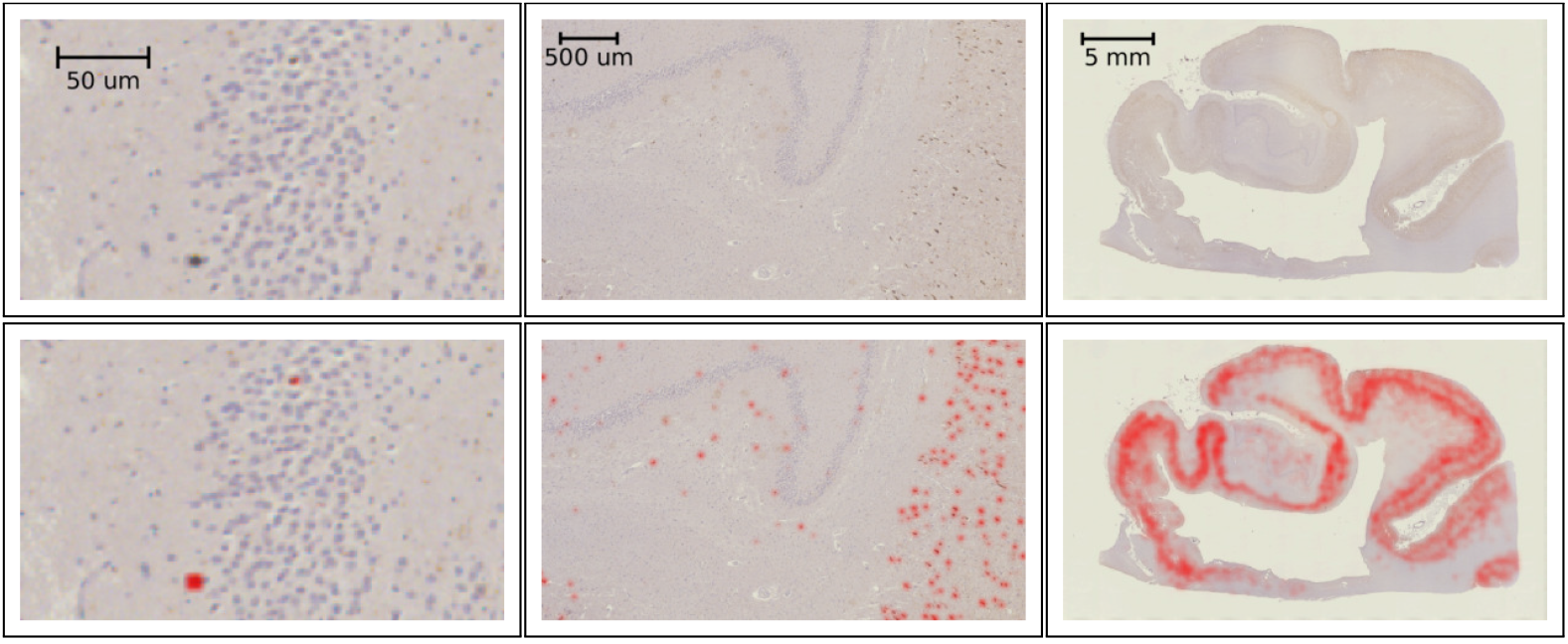
Tau detection results. Tau stained histology sections (top) with the result of our tangle detection (bottom, red) at 3 different scales, zooming out from single tangles (left) to the entire medial temporal lobe (right).

Figure 9 shows several sections of the Mai-atlas along the rostral to caudal axis in millimeters. In the same coordinate system, we show our post mortem MRI with manual segmentations superimposed, and our histology stains and estimated tangle density. For visualization of this sparse data in 3D, interpolation was applied between slices. To sample at a fraction *p* between slices *I* and *J*, a symmetric LDDMM transformation was computed (as in ANTs SyN [47]), and a weighted average of images was computed from the flow: 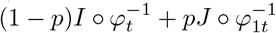.

Finally, Fig. 10 shows our estimated area covered by tau tangles on each page of the Mai atlas for several structures (entorhinal cortex, subiculum, CA fields). We observe a trend of decreasing tangle concentration in the rostral to caudal direction, which will need to be verified for reproducibility as more specimens become available.

**Fig 10.**
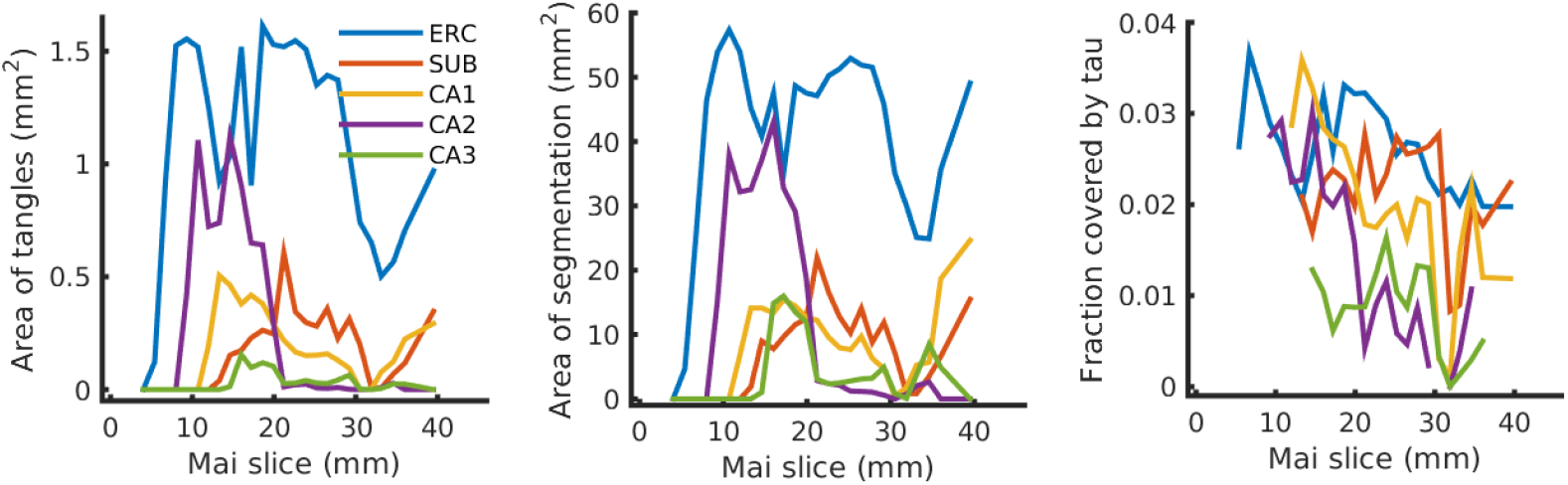
Tau tangles on each page of Mai atlas for several structures. Left shows the total area of tau tangleps detected within several anatomical structures (ERC: entorhinal cortex, SUB: subiculum, CA: Cornu Ammonis). Center shows the area of these structures, and right shows the fraction of this area covered by tau tangles.

## Discussion

In this work we proposed a new image mapping method that accommodates contrast differences, missing data, and artifacts. This was achieved by formulating the imaging process as (i) an unknown shape through the action of the diffeomorphism group, (ii) an unknown change in contrast through the action of polynomial maps, (iii) the addition of Gaussian noise. Here (i) describes the object being imaged, and (ii-iii) describe the imaging process, reflecting the distinction made by Shannon between source and channel. This model allows us to formulate multi modality image matching as a penalized likelihood problem, rather than simply the maximization of an ad hoc image similarity function. This statistical model leads naturally to the formulation of an expectation maximization algorithm that handles missing data or artifacts. We applied this algorithm to simulated images, illustrating its effectiveness for accurate mapping and classification of image pixels, and its superiority over alternatives. For 2D histology we demonstrated the effectiveness of each of our two contributions, contrast mapping and tissue classification, in isolation and simultaneously. Finally we applied this technique to the challenge of reconstructing 3D volumes from histology by mapping to post mortem MRI. In conjunction with convolutional neural networks, this allowed us to map out the 3D distribution of tau tangles in the medial temporal lobe.

Here we demonstrated that for 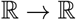 affine contrast transformations, our formulation is equivalent to normalized cross correlation. Another popular image similarity term is mutual information, which is invariant to all invertible transformations. Our approach can accommodate these invariances if we allow for arbitrary nonparametric transformations, which can be thought of as high degree polynomials, or as linear combinations of narrow kernel functions. In this limit, a standard result of statistical prediction results in an intensity transformation given by conditional expectation, *F*(*i*) = *E*_*J|Ĩ*=*i*_[*J*]. This transformation results in a cost function with the same set of invariances as mutual information.

A third popular image similarity term, introduced in [47], is local normalized cross correlation. We are currently extending our method to include polynomial contrast transformations where coefficients are smooth functions of space. As in local normalized cross correlation, this will allow accommodation of image nonuniformity due to magnetic field inhomogeneities or coil sensitivity in MRI, or variable illumination optical imagery.

Typically image registration has involved the balance between a regularization term and a data attachment term in optimization, which is characterized by a single parameter chosen to reflect the researcher’s priorities. A limitation of our algorithm is that it requires more parameters: a variance for shape change (regularization), and variance of image noise, background noise, and artifact noise. These must be chosen carefully to reflect physical characteristics of our imaging model. Further, as is typical of Expectation Maximization algorithms, optimization in our setting can be slow and sensitive to initialization. The choice of polynomials here to describe intensity changes was for simplicity only, linear combinations of any basis functions can be included in the same formulation. The estimation of polynomial coefficients can be unstable, and future work will involve investigating alternative bases.

Our contribution to AD understanding stems from the need to bridge the gap between 3D imaging such as MRI which can be obtained in living subjects over time, and 2D histopathology which is the technique used to make the diagnosis postmortem. While some authors have successfully registered histology to MRI in well controlled conditions [48], we believe that the generative model proposed here, which accommodates variable contrast and missing data, will be a valuable approach for handling typical data moving forward. We are continuing to acquire post mortem samples, and the single subject results presented here will be augmented over the next several years, enabling a more detailed examination of tau distribution. This work advances the field of brain mapping in two important ways. First it moves away from ad hoc image similarities, and toward statistical models of image formation. While our work used a simple white noise model, this framework has the potential to connect with imaging physics and benefit from known properties of imaging systems, such as their signal transfer and noise performance. Second, this method accommodates mapping between images taking values in arbitrary dimensions, in the presence of missing tissue and artifacts. This allows accurate brain mapping to expand from well controlled clinical imaging, to the massive diversity of neuroscience data. For example, in the mouse community, accurate image mapping between Nissl stained tissue and microscopy with multiple fluorophores is commonly required in the presence of variably dissected or damaged tissue. We are currently applying these techniques to CLARITY [19, 20] and iDISCO [49] images in mouse and rat [50], serially sectioned mouse as part of the BRAIN Initiative Cell Census Network [51], and revisiting older datasets where images were excluded due to artifacts or damaged tissue.

## Acknowledgments

This work was supported by the National Institutes of Health (NIH) (www.nih.gov) grants P41EB015909 (SM), R01NS086888 (SM), R01EB020062 (MM), R01NS102670 (MM), U19AG033655 (MM), R01MH105660 (MM); the National Science Foundation (NSF) (www.nsf.gov) 16-569 NeuroNex contract 1707298 (MM); the Computational Anatomy Science Gateway (DT, MM) as part of the Extreme Science and Engineering Discovery Environment (XSEDE [52]) which is supported by the NSF grant ACI1548562; the Johns Hopkins University Alzheimer’s Disease Research Center with NIH grant P50AG05146 (MA); the Dana Foundation’s (www.dana.org) clinical neuroscience research program (MA); the BrightFocus Foundation (www.brightfocus.org) (JT); and the Kavli Neuroscience Discovery Institute (kavlijhu.org) supported by the Kavli Foundation (www.kavlifoundation.org) (DT, MM, JT). The funders had no role in study design, data collection and analysis, decision to publish, or preparation of the manuscript.

